# Coil and flow diverting stents as drug delivery platforms for cerebral aneurysm treatment

**DOI:** 10.1101/2025.06.24.661344

**Authors:** John W. Thompson, Jennifer Suon, Maxon V. Knott, Gina Corsaletti, Naser Hamad, Joshua Hanna, Pedro Barkevitch Rodrigues, Sai Sanikommu, Helena Hernandez-Cuervo, Rianna Haniff, Shunsuke Hataoka, Stephanie H. Chen, Ahmed Abdelsalam, Jayro Toledo, Tiffany Eatz, Sanjoy K. Bhattacharya, Joshua M. Hare, Michal Toborek, Roberto I. Vazquez-Padron, Joshua D. Burks, Evan M. Luther, Robert M. Starke

**Author notes:** **Corresponding Author** John W. Thompson.

## Abstract

Cerebral aneurysm occlusion with coils and flow diverting stents has become the first line treatment for both unruptured and ruptured cerebral aneurysms. As these technologies have advanced, there have been changes in device shape and surface coating to enhance aneurysm embolization while reducing stent thrombogenicity. Drug eluting stents have been used with great success in the targeted delivery of rapamycin, a mTOR complex 1 inhibitor to prevent restenosis in coronary and peripheral artery disease. However, few studies have investigated the use of coils and stents as delivery platforms for sustained drug release to cerebral aneurysm tissue. In this study, we used the bio-compatible and degradable polymers, gelatin and PLGA and a simple evaporative coating technique to investigate the release of rapamycin over time from coated platinum coils and Pipeline flow diverting stents. Rapamycin coated coils were incubated with human vascular endothelial cells *in vitro* to confirm therapeutic levels of rapamycin release. The rate of rapamycin release was similar in both gelatin and PLGA coated coils and was sustained for more than three weeks. Rapamycin was bioactive, at a therapeutic dose and inhibited mTOR complex 1 in human brain endothelial cells treated with a rapamycin coated coil. The relative degree of mTOR complex 1 inhibition was greater in PLGA compared to gelatin coated coils. Coating flow diverting stents with a rapamycin-PLGA coating demonstrated continuous rapamycin release over a 35 day period. Reducing the percent PLGA polymer concentration caused a robust and sustainable release of rapamycin. The PLGA coating was resilient enough to allow device recapturing without affecting rapamycin eluting rates or device deployment and expansion. This work provides a simple, feasible and tunable method to coat occlusion devices for preclinical studies investigating targeted drug delivery for improved parent vessel healing and aneurysm obliteration.

## Introduction

Since the advent of the detachable coils, there have been steady incremental improvements and developments in endovascular aneurysm occlusion devices and techniques, including micro balloons, flow diverters, Woven EndoBridge (WEB), and second-generation coils. ^1–5^ As a result, endovascular intervention has largely become the first line treatment modality for both ruptured and unruptured cerebral aneurysm. However, aneurysm recanalization and retreatment remain a significant risk for patients treated with coil embolization and flow diversion. ^6–8^ Additionally, the risk of thromboembolic events, dual antiplatelet therapy hemorrhagic complications and aneurysm rupture remain following flow diversion stent treatment as the aneurysm is slowly occluded over a period of months. ^9–11^

In an attempt to improve aneurysm occlusion rates, coils have been modified with the addition of gel polymers, growth factors, cytokines, hydrogels, and cells. ^12, 13^ Similar, surface modifications of flow diverting stents have been developed to reduce thrombogenicity while increasing cell attachment and coverage. ^14–16^ In cardiology, drug eluting stents have become a mainstay in coronary artery disease treatment to prevent restenosis. However, there have been limited studies in which coils and flow diverting stents have been used as a platform for drug delivery to cerebral aneurysm tissue for improved vessel healing and aneurysm occlusion.

Extensive research has examined the use of biodegradable polymers as a carrier for device coatings. Some of these polymers include poly (L-lactic acid), poly (D, L-lactic acid), poly (lactic-co-glycolic acid) (PLGA), polycaprolactone, and gelatin to name a few. ^17–19^ These polymers are widely used in the medical field and in preclinical studies to allow for modulation of the drug delivery rates. Several techniques are used for applying a polymer coating which include the use of specialized equipment for ultrasonic spray atomization, spin and melting techniques but also include the simple technique of solvent evaporation.^17, 20^ Therefore, the objective of this study was to determine if solvent evaporation coating (i.e. dip coating) of standard platinum aneurysm coils and Pipeline flow diverting stents could be used as a delivery platform for continuous and sustainable drug release. For this study we used rapamycin as the coating drug since its application has been well characterized within the cardiac field.

## Materials and Methods

### Materials

RPMI-1640 media, Fetal Bovine Serum (FBS), MEM amino acids, sodium pyruvate, and penicillin/streptomycin were purchased from Gibco/Life Technologies (Grand Island, NY). Phospho-AKT (S473), phosphor-S6 ribosomal protein (S235/236), and β-actin were purchased from Cell Signaling Technology (Danvers, MA). NuSerum, Bicinchoninic acid (BCA), secondary antibodies and enhanced chemiluminescence (ECL) were purchased from ThermoFisher Scientific (Waltham, MA). All other reagents were purchased from Sigma-Aldrich (St. Louis, MO) unless otherwise noted.

### Coating stents and coils with rapamycin

Standard platinum aneurysm coils (Penumbra Inc; Alameda, CA) and Pipeline flow-diverting stents (Medtronic; Minneapolis, MN) were used for this study. A PLGA polymer coating solution was created consisting of 1% or 0.3% 50:50 poly-D,L-lactic glycolic acid (PLGA), magnesium hydroxide, and 50 μM or 250 μM rapamycin in dichloromethane anhydrous. A magnesium hydroxide suspension was initially prepared in dichloromethane anhydrous and sonicated at room temperature for 10 minutes before adding it to the PLGA solution at a 20% wt/vol ratio of PLGA. ^17^ Coils and stents were coated via immersion in the PLGA coating solution with gentle mixing for 20 seconds. The devices were then removed and allowed to dry at room temperature for 10 minutes before being stored overnight at 4°C in the dark. In some experiments, coils were coated by dipping in a 1% gelatin solution in phosphate buffered saline (PBS) containing 250 μM rapamycin either alone or on top of an already applied PLGA-rapamycin coating and stored as above. Control coating solutions and devices were prepared and treated as above but without the addition of rapamycin.

### Rapamycin elution assay

To determine the release profile of rapamycin, the coils and stents were placed in 1.5 mls of PBS and incubated at 37°C with gentle rocking. At time intervals, the coils and stents were removed and placed into fresh 1.5 mls PBS. Rapamycin release into the PBS was determined using a NanoDrop 2000 UM-Vis spectrophotometer (Thermo Fisher; Waltham, MA) at 277nm. The nanodrop was calibrated using a rapamycin curve prior to each day’s measurements.

### Cell culture and western blot analysis

Human brain microvascular endothelial cells (ScienCell Research Lab; Carlsbad, CA) were cultured in RPMI-1640 media supplemented with 10% FBS, 10% NuSerum, 1% MEM amino acids, 1% sodium pyruvate, and penicillin/streptomycin. For experimentation purposes, the cells were plated in 60mm dishes and grown to confluency prior to treatment. For western blot analysis, the cells were lysed in RIPA Buffer [20 mM Tris-HCl (pH 7.5), 150 mM NaCl, 1 mM EDTA, 1% NP-40, 1% sodium deoxycholate, 2.5 mM sodium pyrophosphate, 1 mM Na_3_VO_4_ and 1 mM PMSF]. Protein concentration was determined by bicinchoninic acid (BCA), and 15 ug of protein was loaded onto a 12% SDS-polyacrylamide gel and electroblotted to nitrocellulose.

Membranes were blocked in 5% dry milk/tris-buffered saline with 0.1% Tween 20 detergent (TBST) and hybridized with primary antibodies overnight at 4°C. Blots were probed with phosphorylated AKT (S473), phosphorylated S6 ribosomal protein (S235/236), and β-actin. Membranes were washed with TBST followed by incubation with secondary antibodies for 1 hr at room temperature. Proteins were detected using enhanced chemiluminescence (ECL) system.

### Statistics

All data are expressed as mean ± S.E.M. Statistical analysis between two groups was carried out using the unpaired Student’s t-test. Statistical analysis between more than two groups was performed using a one-way ANOVA with Dunnett’s multiple comparison post hoc test. A p-value less than 0.05 was considered statistically significant for all analyses.

## Results

A rapamycin concentration curve ranging from 3.0 to 100ug/ml was prepared in PBS and 1ul of the solution was analyzed using a NanoDrop UV spectrophotometer at wavelength 277nm. As shown in Figure 1, the lower sensitivity threshold for rapamycin detection was 6.25 ± 0.58ug/ml with a correlation coefficient of 0.9933, demonstrating both excellent linearity and reproducibility (Figure 1).

**Figure 1:**
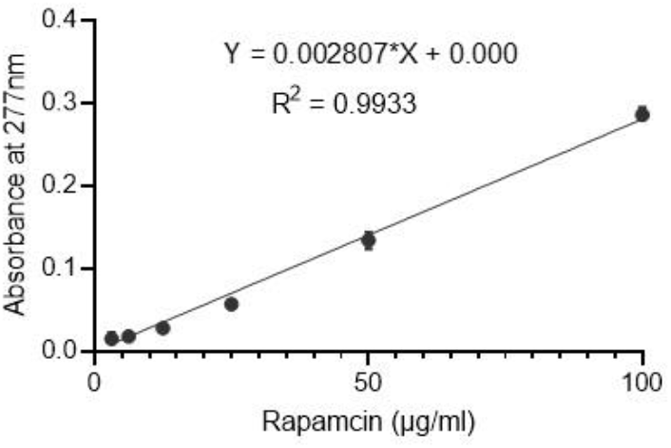
Rapamycin calibration curves using a NanoDrop 2000 UM-Vis spectrophotometer at 277nm. N = 4, SEM.

Uncoated coils have a void space created by the spiraling platinum coil material. We found that coating the coil with both gelatin and PLGA filled this dead space with the coating polymer (Figure 2a). Unlike gelatin though, the PLGA polymer was also found to completely cover the coil forming a layer over the coil which was macroscopically visible as a rough white coating (Figure 2a).

**Figure 2:**
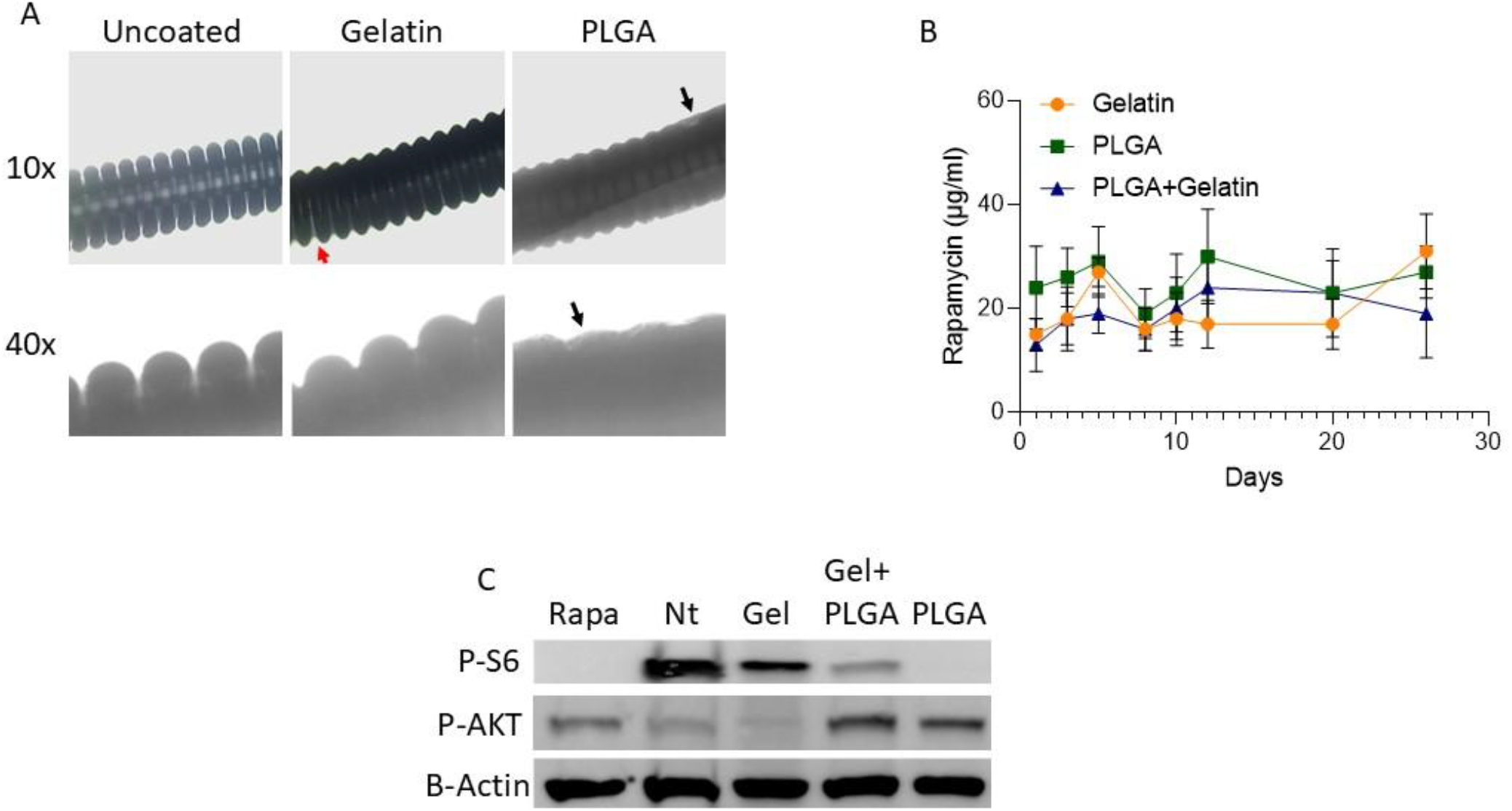
Coating and testing of rapamycin-releasing coils using 1% gelatin, 1% PLGA, and 1% gelatin-1%PLGA. A. Bright field microscopy demonstrating coil groove filling by gelatin and PLGA. Red arrow points to visible gelatin within a groove; black arrow points to surface coating by PLGA. B. Sustained release of rapamycin over a 26-day period from gelatin, PLGA, and gelatin-PLGA-coated coils. C. Human vascular endothelial cells treated with rapamycin coated-gelatin (Gel), gelatin-PLGA (Gel-PLGA), and PLGA-coated coils for 24 hrs. The ability of the eluted rapamycin to inhibit mTOR complex 1 was determined be the phosphorylation of downstream indicator protein Ribosomal S6 protein, while mTOR complex 2 was determined by changes in AKT (S473) phosphorylation. N = 3, SEM.

Next, we determined the release of rapamycin from gelatin, PLGA, and gelatin-PLGA coatings. As shown in Figure 2B, rapamycin release from PLGA, gelatin, and gelatin-PLGA coatings exhibited similar rapamycin release rates. Rapamycin release was detectable within 24 hours and was found to continue to be released for at least 26 days in all three of the coating polymers. The multiday release average was similar for each coating polymer: gelatin alone (20 ± 2.1 ug/ml), gelatin-PLGA (19 ± 1.3 ug/ml), and PLGA alone (24.9 ± 1.2 ug/ml).

To confirm that the coating polymer and solvent did not alter rapamycin biological activity, gelatin, PLGA, and gelatin-PLGA coated coils which had been incubated in PBS for 28 days were placed on top of a confluent layer of human brain microvascular endothelial cells for 24hrs. The cells were then harvested for western blot analysis, and mTORC1 and mTORC2 activity determined by changes in phosphorylation of the known mTOR activation proteins AKT and ribosomal protein S6.^21, 22^ As shown in Figure 1c, phosphorylated S6 ribosomal protein, an indirect downstream mTORC1 target, was completely inhibited by rapamycin eluting from PLGA coils and was similar to that observed in cells treated directly with rapamycin. In contrast, S6 ribosomal protein phosphorylation was substantially reduced but not eliminated in cells treated with rapamycin eluting from either gelatin alone or gelatin-PLGA coatings. We also observed a parallel increase in mTORC2 activation as evidenced by increased AKT S473 phosphorylation in rapamycin treated cultures and cultures treated with PLGA coils. Interestingly, AKT phosphorylation was similar in gelatin-PLGA treated cultures compared to PLGA alone. These results indicate that the coating solutions do not affect rapamycin function and that the eluting rapamycin, even after prolonged periods, is at a therapeutic concentration.

Based upon our coil results demonstrating increased mTORC1 inhibition with the PLGA polymer, we decided to test the PLGA polymer only for stent coating. Similar to coils, PLGA evaporation coating of flow diverting stents formed a white coating on the stent surface. Microscopic imaging revealed the PLGA polymer formed small aggregates on the struts of the stents (Figure 3a). Next, we determined rapamycin release from the coated stent. As shown in Figure 3b, rapamycin was detectable at 15ug/ml at day 5 and was found to be continuously released at similar concentrations over a 35 day period with an average multiday release of 20 ± 1.6 ug/ml.

**Figure 3:**
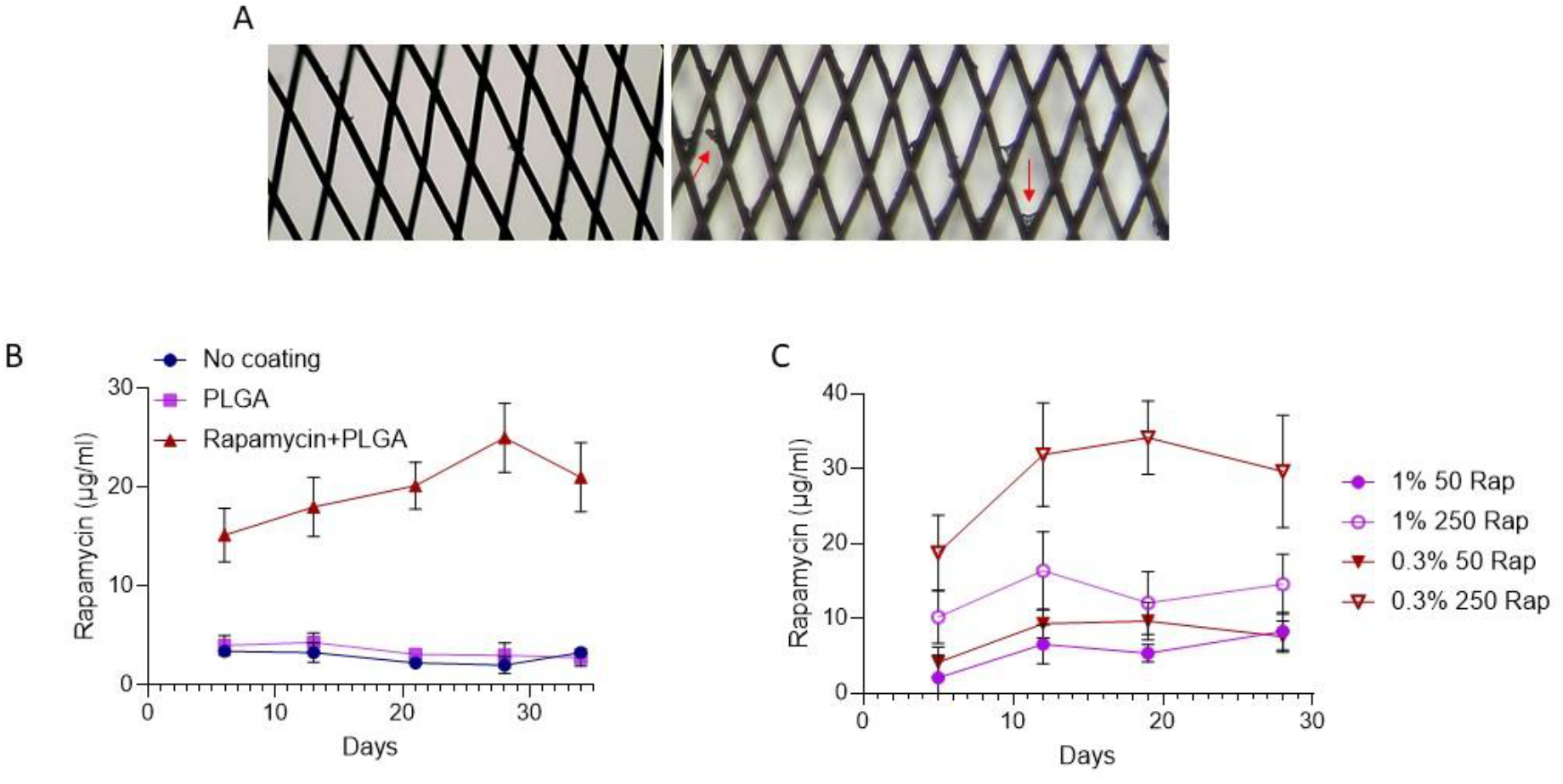
Coating and testing of rapamycin-releasing flow diverters using PLGA. A. Micrograph of control and 1% PLGA-coated flow diverting stent. B. Sustained release of rapamycin over a 35-day period. C. Rapamycin release overtime from stents coated with 50uM or 250uM Rapamycin (Rap) and 0.3% and 1% PLGA. N = 4, SEM.

We then determined if rapamycin elution was tunable by altering the percentage of PLGA polymer and the concentration of rapamycin in the coating polymer. As shown in Figure 3c, the rate of rapamycin release was significantly increased when coated with a 0.3% PLGA solution compared to a 1% PLGA solution, which nearly doubled the multiday accumulation averages 28 ± 2.6 and 13 ± 0.66, respectively. As would be expected, reducing the concentration of rapamycin in the coating solution from 250 μM to 50 μM resulted in a corresponding reduction in rapamycin release concentrations.

To coat a flow diverting stent or coil for preclinical aneurysm occlusion studies, the stent or coil would have to be recaptured following polymer-drug coating. Therefore, we tested if the PLGA coating impeded stent recapture and if the PLGA-rapamycin coating was affected by stent recapturing. To test this, flow diverting stents were maximally deployed and dipped in a rapamycin-PLGA polymer coating solution and allowed to dry overnight. The stent was then recaptured and partially deployed 2 times before it was placed in 1.5 mls PBS solution for 10 days. As shown in Figure 4a, rapamycin elution from recaptured stents was not significantly different from rapamycin-PLGA control stents which were not recaptured.

**Figure 4:**
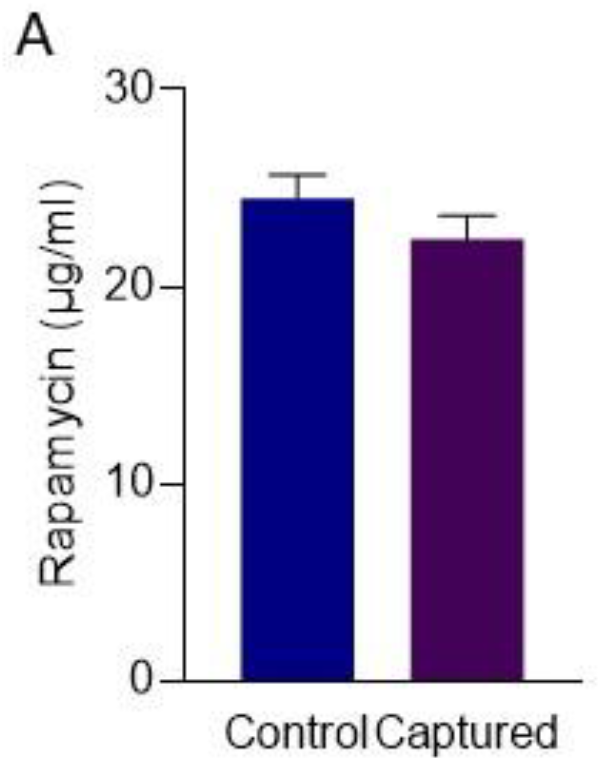
PLGA coating does not affect flow diverting stent function. A. Flow-diverting stents were coated with 250uM rapamycin-1% PLGA and the durability of the PLGA coating determined by capturing-deploying the stent two times. Rapamycin concentration was determined at 10 days and compared to a rapamycin-1% PLGA-coated stent, which was not captured. N = 3. SEM.

## Discussion

In this study, we demonstrate the feasibility of using a simple evaporation technique to coat both flow diverting stents and coils, which can be used in preclinical studies for the targeted delivery of drugs to aneurysm tissue. Our results demonstrate: 1) sustained drug (rapamycin) release for at least 34 days, 2) the eluting drug was therapeutic and able to inhibit mTORC1 activation, 3) the concentration of drug eluting for the device was tunable by either altering the coating drug concentration or by reducing the PLGA coating concentration, and 4) PLGA coating is robust enough to allow device recapturing without affecting device function.

The current literature consists of investigations of PLGA-coated coils but not stents for the treatment of aneurysms. Specifically, PLGA-coated coils have been used for the delivery of MCP-1 and OPN to improve aneurysm healing in mouse aneurysm models.^23^ While in humans, clinical trials have shown that PLGA-coated coils are safe for aneurysm treatment but were ineffective in increasing aneurysm obliteration.^24^ Notably, our study is the first to our knowledge to investigate PLGA coating of flow-diverting stents for drug delivery to aneurysmal sites.

Both gelatin and PLGA polymers are used as scaffold and surface coatings in medical applications. These polymers are characterized by predictable rates of biodegradation and excellent biocompatibility. However, as PLGA is degraded, it generates lactic and glycolytic acid which causes a localized inflammatory reaction in the surrounding tissues. Therefore, magnesium hydroxide was added to the drug-PLGA coating to neutralize the acidic PLGA degradation products by generating Mg2+ and OH- as the magnesium hydroxide is eluted. The addition of magnesium hydroxide to PLGA coating solution has been shown to improve vessel healing in the proximal area of the device. ^17^ Our results demonstrate that the addition of magnesium hydroxide to the coating had no effect on rapamycin biological activity.

There are several techniques used to incorporate drugs onto a delivery platform. These include either a chemical association of the drug to the device’s surface or a coating technique in which a liquid is applied to the surface via either electro-treatment, plasma, spray, or dip coating. Unlike the other methods which require costly special equipment, dip coating is a simple and cost-effective method which can be efficiently performed in any laboratory. Therefore, in this study we used a dip coating technique to coat both platinum coils and flow diverters, primarily composed of cobalt-chromium. The dip coating technique consists of placing the device in a polymer solution which is then removed. The polymer solvent is then allowed to evaporate, thereby encapsulating the drug in the polymer as it resolidifies. The drug is released as the polymer is slowly dissolved in an aqueous solution such as blood. Therefore, as in this study, drug release could be modulated by either decreasing the drug loading concentration or by altering the percent PLGA. Furthermore, by altering the coating and solvent materials used the drip coating technique could be expanded to allow for a “layering” of different drugs which are differentially released as the outer layers are slowly dissolved. ^23^

## Conclusion

This study demonstrates a simple, practical, and adaptable method for the coating of clinically available occlusion devices, which can be used for research on vessel healing through sustained drug delivery to aneurysmal tissue. However, initial experiments need to be conducted to confirm drug release concentrations and that the coating polymers and solutions do not interfere with the activity of the drug to be investigated.

